# Organoid Polymer Functionality and Mode of *Klebsiella Pneumoniae* Membrane Antigen Presentation Regulates Ex Vivo Germinal Center Epigenetics in Young and Aged B Cells

**DOI:** 10.1101/2020.02.16.947408

**Authors:** Pamela L. Graney, Kristine Lai, Sarah Post, Ilana Brito, Jason Cyster, Ankur Singh

## Abstract

Antibiotic-resistant bacteria are a major global health threat that continues to rise due to a lack of effective vaccines. Of concern are *Klebsiella pneumoniae* that fail to induce *in vivo* germinal center B cell responses, which facilitate antibody production to fight infection. Immunotherapies using antibodies targeting antibiotic-resistant bacteria are emerging as promising alternatives, however, cannot be efficiently derived ex vivo, necessitating the need for immune technologies to develop therapeutics. Here, four-arm PEG-organoids were developed to elucidate the effects of polymer end-point chemistry, integrin ligands, and mode of *K. pneumoniae* antigen presentation on germinal center-like B cell epigenetics, to better define the cell-microenvironment factors regulating *ex vivo* germinal center dynamics. Notably, PEG vinyl sulfone or acrylate failed to sustain primary immune cells, but functionalization with maleimide (PEG-4MAL) led to B cell expansion and germinal center-like induction. RNA sequencing analysis of lymph node stromal and germinal center B cells showed niche associated heterogeneity of integrin-related genes. Incorporation of niche-mimicking bioadhesive peptides revealed that collagen 1 mimicking peptides promoted germinal center-like dynamics and epigenetics. PEG-4MAL organoids elucidated the impact of *K. pneumoniae* membrane embedded protein antigen versus soluble antigen presentation on germinal center-like activation and preserved the response across young and aged mice.

## 1. Introduction

Bacterial infections remain a major global health problem, causing more than 5 million deaths annually^1^. Antibiotics are currently the standard medical intervention for several bacterial pathogens, including *Mycobacterium tuberculosis, Clostridioides difficile, Escherichia coli*, *Pseudomonas aeruginosa*, and *Klebsiella pneumoniae*, among others, for which we do not yet have vaccines. Even more concerning are the rising antibiotic resistance threats in the United States and worldwide. According to the 2019 Antibiotic Resistance Threats Report by the U.S. Center for Disease Control^2^, more than 2.8 million antibiotic-resistant infections occur in the U.S. each year, and more than 35,000 people die as a result. In addition, 223,900 cases of *Clostridioides difficile* occurred in 2017 and at least 12,800 people died. According to WHO, in 2016, an estimated 490,000 people worldwide developed multidrug-resistant tuberculosis (MDR-TB), and an additional 110,000 people with rifampicin-resistant TB were also newly eligible for MDR-TB treatment. In recent years, the emergence of broad MDR strains of *K. pneumoniae*, which not only causes pneumonia and infections of the blood stream and urinary tract, but also are not cleared by β-lactam antibiotics, including carbapenem and cephalosporins, or by non-β-lactam antibiotics, such as aminoglycosides and tetracycline, has made *K. pneumoniae* one of the most concerning threats^3^. Vaccines are still unavailable for several bacterial infections, including most antibiotic-resistant bacteria, and vaccines to prevent such infections can only be best developed in a timely manner on the basis of our increasing insights into the immune response. Effective vaccination also requires identification of bacterial antigens, their immunogenicity, and the mode of presentation of these antigens – as embedded bacterial antigens in outer membranes or in soluble purified forms, making successful vaccination challenging. New immunotherapies using monoclonal antibodies targeting bacterial antigens are emerging as promising alternatives to passive immunization to tackle antibiotic-resistant bacteria, including *K. pneumoniae ^4, 5^*, but often fail to mediate broad protection against different serotypes.

Developing antibodies with high cross-reactivity requires enhanced understanding of the underlying immune response, which typically involves a germinal center B cell response in lymph nodes that makes high-affinity antibodies through iterative somatic hyper mutation and affinity maturation processes^6, 7^. Unfortunately, the polysaccharide antigens on many gram-negative bacteria fail to induce robust germinal center responses *in vivo*, which necessitates the development of an *ex vivo* immune technology, where the germinal center reaction can be induced irrespective of antigen type and form. However, the development of such immune technologies is non-trivial as factors that spatially and temporally regulate the *ex vivo* germinal center response, ranging from microenvironment to antigen format, are poorly understood. The goal of the current study is to identify these factors and engineer a materials-based immune technology.

During an immune response to bacterial and viral infections, naïve B cells in the lymph node and spleen, encounter antigen and form sub-anatomical structures within B cell follicles in secondary lymphoid organs, referred to as germinal centers. Antigen-activated germinal center B cells undergo rapid proliferation and somatic hypermutation of their immunoglobulin variable genes. The proliferating B cells mutate their B cell receptors to produce mutant germinal center B cell clones with a range of affinities against the activating antigen^8, 9^, which regulates cross-reactivity against serotypes. The germinal center response is a complex process involving multiple cell types and environmental cues^10–12^. B cell follicles are composed of B cells, CD40 ligand (CD40L) presenting follicular T helper (T_FH_) cells, B cell activating follicular dendritic cells (FDCs)^13–15^, extracellular matrix (ECM) proteins such as Arg-Gly-Asp (RGD)-presenting vitronectin^16, 17^, and several other factors. For example, integrin-mediated interactions between B cells and FDCs influence germinal center B cells^12^. B cell activation is marked by epigenetic regulation and immune receptor signaling, which come into play at critical transitional stages of the germinal center reaction^18, 19^. In the germinal center reaction, there is transient suppression of enhancers and promoters of genes that regulate immune signaling pathways, antigen presentation, and checkpoints, which revert to the active state when germinal center B cells are signaled to exit the germinal center reaction. Exit from the germinal center reaction occurs by differentiating into antibody-producing plasma cells or memory B cells. Currently, the tissue and microenvironment factors governing B cell dynamics, temporal expression of phenotypic and epigenetic markers, and exit from the germinal center reaction are not well understood.

Early efforts to elucidate B cell differentiation involved a controlled environment of naïve B cells co-cultured in 2D with stromal cells, called 40LB. These cells were engineered to provide critical signals for B cell survival and differentiation, including the T cell signal CD40L and B cell activating factor (BAFF), secreted *in vivo* by FDCs. However, the *in vitro* 2D presentation of these signals is not sufficient for induction of a germinal center response that resembles an *in vivo*-like response^19^, suggesting a critical role for the 3D microenvironment. Indeed, we have previously shown that incorporation of naïve B cells and 40LB in 3D organoids engineered from gelatin and silicate nanoparticles can significantly drive germinal center B cell phenotype, transcriptome, and somatic hypermutation just after 4 days *ex vivo*^19^, similar to *in vivo* immunized mice. To better decouple the signals driving this response, we further engineered modular organoids with tunable microenvironments using a four-arm polyethylene glycol presenting maleimide (PEG-4MAL) functionalized with thiolated ECM-mimicking peptides and enzymatically degradable crosslinkers^20^. Using this system, we have shown that interactions between integrins and VCAM-1-mimicking peptide regulated phosphorylation of key proteins in the B cell receptor (BCR) signaling^20^. However, to date, the role of PEG end-group chemistry in regulating the germinal center response and contributions of microenvironment-epigenome interactions have not yet been explored, despite that such interactions would be expected to regulate cell behavior.

In the current work, our goal was to identify factors, including polymer end-point chemistry, integrin ligands, and mode of *Klebsiella Pneumoniae* antigen presentation, in a synthetic immune organoid that regulates the phenotype and epigenetics of *ex vivo* germinal centers and provide a future opportunity to derive age-specific germinal center B cells, and subsequently antibody-secreting cells and antibodies against a large set of bacterial antigens that can be injected into patients as therapeutics. We provide the first comparison of advanced PEG-4MAL hydrogels with those functionalized with vinyl sulfone and acrylate in regulating the fate of primary B cells. We provide a rational methodology to develop immune organoids, driven by single cell RNA sequencing of uniquely identified lymph node stromal cells and bulk RNA sequencing of germinal center B cells from immunized mice to define microenvironment factors to be tested for germinal center induction. We further elucidate the mechanisms by which PEG-4MAL facilitates the germinal center reaction by comparing the time dependent effects of integrin-binding peptides on regulating the kinetics of B cell EZH2, H3k27Me3 epigenetics and BCL6 expression. Subsequently, for the first time to the best of our knowledge, we demonstrate the application of PEG-4MAL as a platform to probe the potential for inducing germinal center responses and regulating epigenetics with bacterial membrane antigens from a clinical *K. Pneumoniae* isolate in their soluble form or membrane embedded form. Finally, the current study sets out to demonstrate the effect of *K. Pneumoniae* antigens in regulating the germinal center phenotype and histone modification in B cells from aged mice >2 years old. These findings describe, for the first time, a direct comparison of PEG end-point chemistry in the engineering of immune organoids and highlight the potential of synthetic germinal center-like organoids as a platform for future development of antigen- and age-specific antibodies for therapeutic intervention, as well developing therapeutics for aged individuals.

## 2. Results and Discussion

### 2.1. RNA sequencing of mouse lymph node stromal cells and germinal center B cells reveals expression of cell-cell and cell-ECM binding genes

Stromal cells establish the compartmentalization of lymphoid tissues, which are critical to the immune response (Figure 1A). Recently, we reported single-cell RNA sequencing of lymph node stromal cells using barcoded droplets (Figure 1A) that revealed nine peripheral lymph node non-endothelial stromal cell clusters^*15*^. Included were the established subsets, C-C Motif Chemokine Ligand 19 (Ccl19)^hi^ T-zone reticular cells (TRCs), marginal reticular cells (MRCs), and follicular dendritic cells (FDCs). The study further identified Ccl19^lo^ TRCs, C-X-C Motif Chemokine Ligand 9 (Cxcl9+) TRCs in the T-zone and interfollicular region, and clusters corresponding to medullary, perivascular and capsular stromal cells. In these studies, a droplet-based single cell RNA sequencing was performed on isolated peripheral lymph node CD45-CD31-cells from mice immunized intravenously with 2.5 × 10^5^ pfu of Lymphocytic choriomeningitis virus (LCMV)-Armstrong infected mice (day 15 post-infection)^*15*^, thereafter performing sequencing and analysis on > 14,000 single cells.

**Figure 1:**
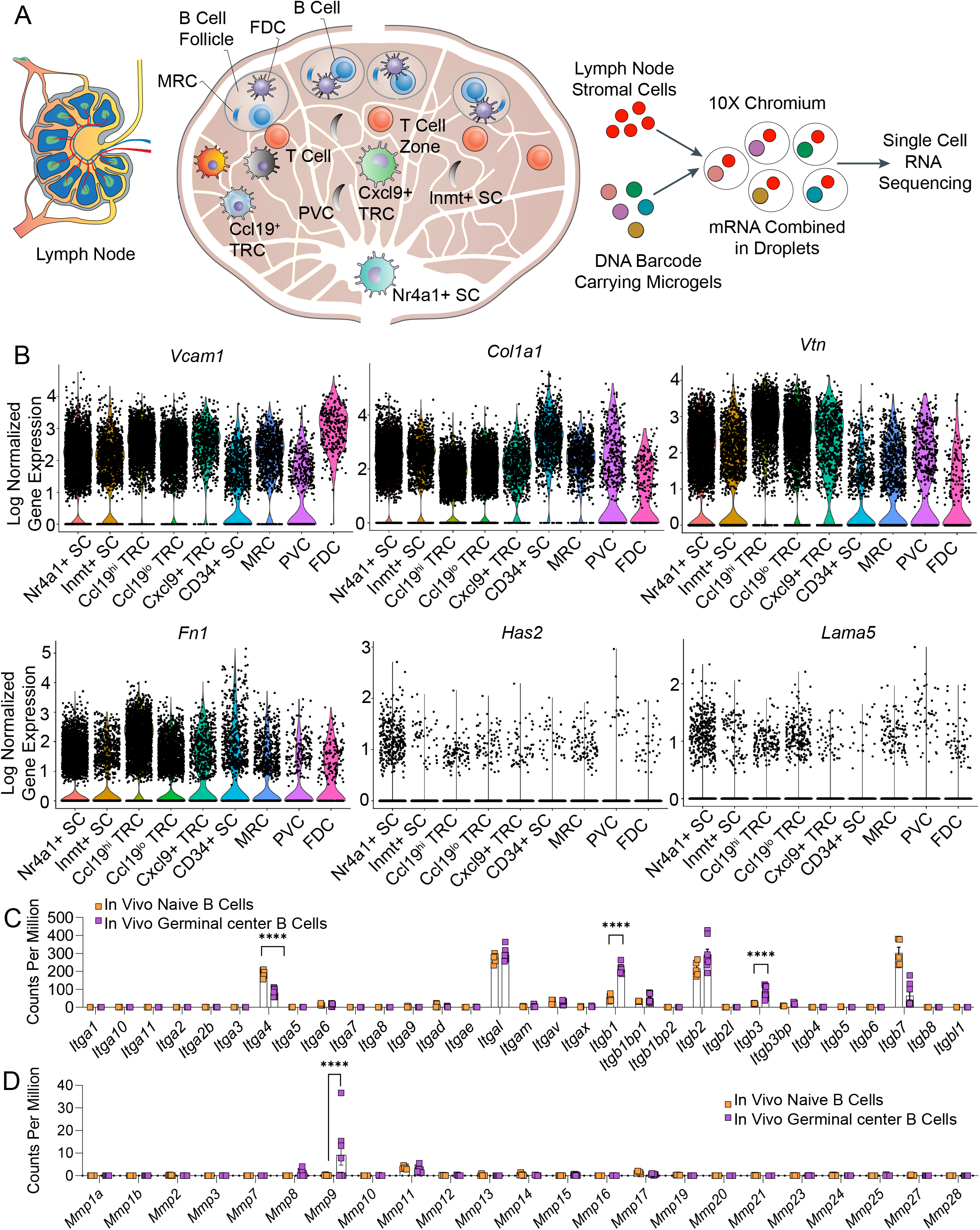
RNA sequencing of lymph node stromal and germinal center B cells shows niche associated heterogeneity of integrin-related genes. **A)** Schematic of lymph node structure comprised of distinct stromal cell populations inside and outside of B cell follicle. Non-endothelial stromal cells isolated from the peripheral lymph node are subjected to droplet-based single cell RNA sequencing workflow. **B)** Single-cell RNA sequencing of nine uniquely identified clusters of peripheral lymph node non-endothelial stromal cells reveals expression of mRNA encoding for proteins that bind to integrins. Violin plots represents the log-normalized gene expression (unique molecular identifier counts) for each cell grouped by cluster. **C)** RNA sequencing comparison of in vivo integrin expression between naïve and germinal center B cells. **D)** RNA sequencing comparison of in vivo matrix metalloproteinase (MMP) expression between naïve and germinal center B cells. In C-D, data represent mean ± S.E.M. Significant differences determined using a 2-way ANOVA with Sidak’s multiple comparison test. ****p<0.0001; N = 5-8.

We analyzed this single-cell RNA sequencing dataset of lymph node stromal cells, including cells located in the B cell follicle, to determine various genes that bind to integrins. We observed that a large fraction of FDCs and MRCs expressed high levels of collagen 1 (*Col1a1*), vascular cell adhesion molecule 1 (*Vcam1*), vitronectin (*Vtn*), and fibronectin (*Fn1*) genes (Figure 1B). VCAM1 protein binds to integrin α4β1 on B cells and is involved in B cell activation, however the role of collagen 1 that binds to α1β1, α2β1, α10β1 and α11β1 integrins, and α5β1 and αvβ3-binding fibronectin and vitronectin are less known. In contrast, laminin (*Lama5*) and hyaluronan (*Has2*) were not abundant in these stromal cells (Figure 1B). We further performed RNA sequencing analysis on naïve versus germinal center B cells isolated from immunized mice, as reported in our recent study^19^, and observed that β1 and β3 were significantly upregulated in *in vivo* germinal center B cells compared to primary naïve B cells (Figure 1C). Integrins containing the β1 subunit can engage with collagen, fibronectin, VCAM-1, and several other matrices. Integrins α4, which forms a heterodimer with β1 subunit, can bind to fibronectin and VCAM-1, and were expressed in both naïve and germinal center B cells. In addition, we also observed high expression of αL integrin subunit, which forms a heterodimer with β2 subunit and binds to ICAM-1 (Intercellular Adhesion Molecule 1). These findings motivate the need to investigate hydrogel networks that can present bio-adhesive ligands of interest, such as collagen 1 and VCAM-1 mimics, to understand the role played by various cell-cell and cell-matrix binding proteins on germinal center phenotype and epigenetics. Another important observation from the germinal center B cell RNA sequencing was upregulation of matrix metalloproteinase (MMP)-9 compared to naïve B cells (Figure 1D). This observation supports the need for developing immune organoids that can be remodeled by MMP-9 secreted by activated B cells, as demonstrated in the next section.

### 2.2. The end group functionality in PEG hydrogels impacts B cell survival and induction of germinal center response

To study the effect of various extracellular matrices and integrin binding proteins found in the B cell follicle, we chose to use Poly(ethylene glycol) (PEG) hydrogels based on 4-arm PEG macromers with either terminal vinyl sulfone (PEG-4VS), acrylate (PEG-4Acr), or maleimide groups (PEG-4MAL) (Figure 2A). These polymers can be functionalized to present integrin-specific peptides that are cysteine-terminated via a Michael-type addition reaction and crosslinked using di-thiolated crosslinkers at physiological pH and temperature under mild conditions^21^, as opposed to UV crosslinking (Figure 2B). While all three chemical functionalities in PEG hydrogels have been studied for a wide range of mammalian cells, they have never been compared directly to each other for their impact on primary B cells for the induction of germinal centers. This is an important biomaterial consideration given the growing interest in building immune tissues^22–24^ and inability of natural matrices in supporting the diverse set of immune cells *ex vivo*.

**Figure 2:**
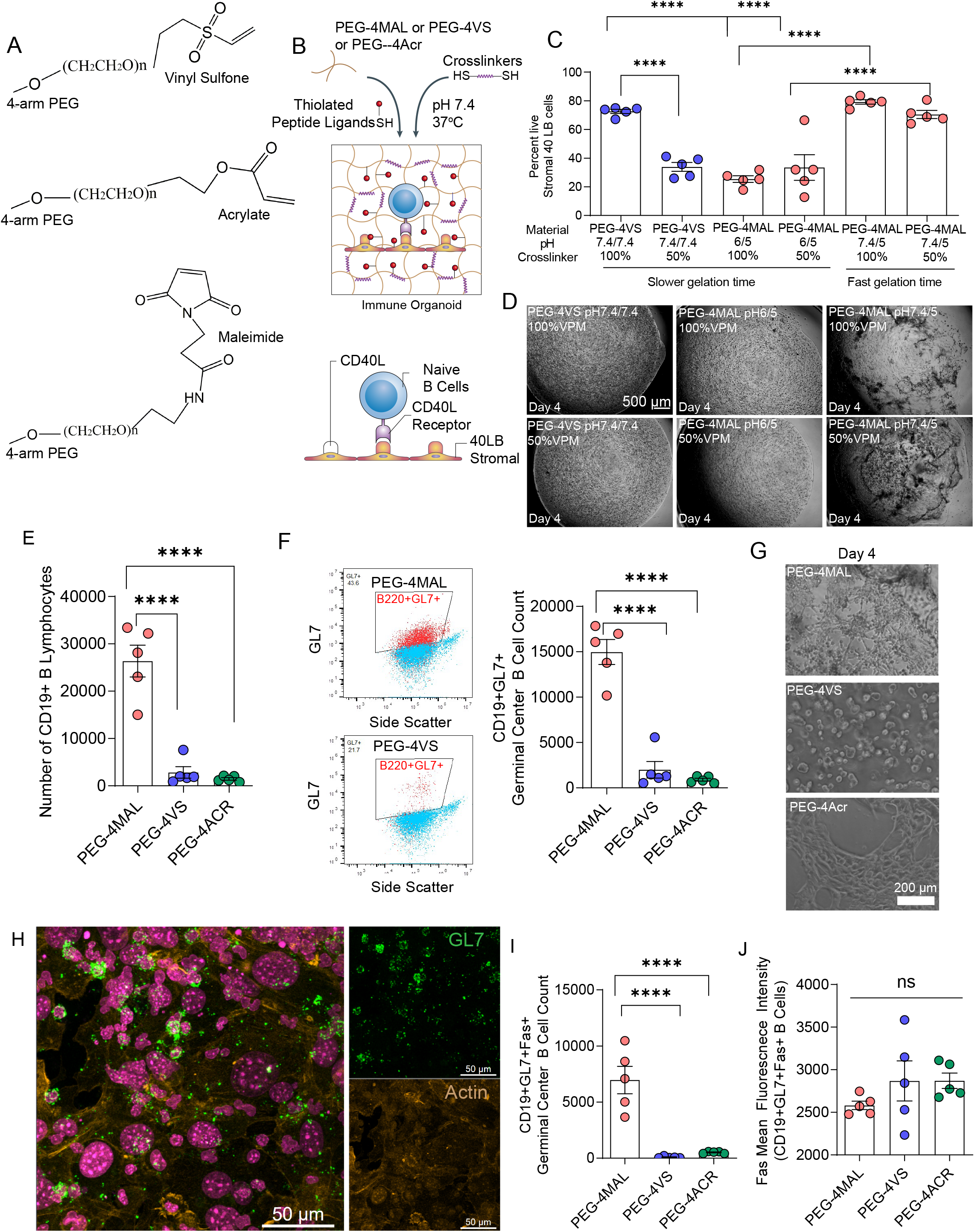
Hydrogel macromer end-modification chemistry regulates survival and differentiation of primary immune cells. **A)** Chemical structure of the vinyl sulfone, acrylate, and maleimide functional groups at the end of four-arm PEG. **B)** Schematic of immune organoid formation with co-cultured CD40L- and BAFF-presenting stromal cells and primary, naive B cells. **C)** Effects of pH and degradability on 40LB cell viability in PEG-4VS and PEG-4MAL hydrogels. **D)** Representative brightfield images illustrating the effects of pH and degradability on stromal cell network formation in PEG-4VS and PEG-4MAL organoids. **E)** Quantitative flow cytometric analysis of CD19+ B cell in four-arm PEG hydrogels modified with maleimide, vinyl sulfone, or acrylate, indicating survival in hydrogels. **F)** Flow cytometric gating strategy for germinal center-like B cell (CD19+GL7) formation in four-arm PEG hydrogels modified with maleimide and vinyl sulfone. **G)** Effects of vinyl sulfone and acrylate functionalization on stromal cell network formation and survival of B cells in PEG-based organoids. **H)** Confocal imaging of germinal center-like B cell (GL7+) and its relative localization with 40LB stromal in PEG-4MAL organoids. Green: GL7, Yellow: Actin, Magenta: DAPI. **I)** Quantitative flow cytometric analysis of proliferative germinal center-like B cell (CD19+GL7+Fas+) formation in four-arm PEG hydrogels modified with maleimide, vinyl sulfone, or acrylate. **J)** Effect of four-arm PEG end-modification chemistry on Fas expression in CD19 +GL7+ B cells. All data represent mean ± S.E.M. and were analyzed via one-way ANOVA with Tukey’s post-hoc multiple comparisons test. ****p<0.0001, ns=not significant; N=5.

We first characterized the survival of our engineered stromal cell 40LB^25, 26^ (developed and provided by Daisuke Kitamura^27^), which are BALB/c 3T3 fibroblasts stably transduced with both T cell-derived CD40 ligand (CD40L) and FDC-secreted BAFF. We tested conditions where the hydrogel macromers formed fast gels (< 1min) versus slow gels (>1 min) by tuning the pH of the PEG macromer and crosslinker type. Specifically, we tested the impact of the ratio of MMP-9 degradable crosslinker peptide (GCRD**VPM** ↓ SMRGGDRCG, referred as VPM) with non-degradable crosslinker dithiothreitol (DTT). The rationale for using VPM was that our RNA sequencing has revealed significantly higher expression of MMP-9 in germinal center B cells as compared to naïve B cells, but other MMPs were indifferent (Figure 1D). We observed high viability among fast gelling PEG-4MAL hydrogels irrespective of the crosslinker pH and VPM:DTT ratio (50 or 100% VPM) (Figure 2C). In contrast, PEG-4VS at 50% VPM or PEG-4MAL at either 50% or 100% VPM, both of which gelled slowly at lower PEG macromer (pH 6), showed cytotoxic effects on 40LB stromal cells (Figure 2C). There were no major differences between the stability of hydrogels (Figure 2D), except that slow degrading hydrogels formed a more uniform hydrogel than fast gelling ones, which can be attributed to wrinkling due to rapid pipetting of fast gelling hydrogels. We, therefore, continued with formulations that showed high viability so that the effects can be delineated between B cells and 40LBs.

We next examined the effect of polymer chemistry on B cell viability and induction of germinal center by co-encapsulating freshly isolated naïve B cells and 40LBs in the hydrogel organoid and culturing in the presence of 10 ng/mL IL-4, regular RPMI with 10% FBS and 1% antibiotics. In these initial studies, we chose a VCAM-1 mimicking peptide REDV, recently reported by us^20^. Interestingly, unlike 40LB cells, primary B cells failed to survive and proliferate in both vinyl sulfone and acrylate functionalized organoids (Figure 2E, F). In contrast, B cells cocultured in PEG-4MAL organoids yielded significantly higher number of CD19+ B cells and CD19+GL7+ B cells (Figure 2E, F, G). The GL7 expressing B cells (green) co-localized with 40LB stromal cells (yellow, spread), as revealed by confocal imaging studies (Figure 2H). Collectively, this is the first study, to the best of our knowledge, that establishes the superiority of advanced PEG-4MAL hydrogels with those functionalized with vinyl sulfone and acrylate in regulating the fate of primary B cells towards induction of germinal centers.

To further understand the plausible reason behind differences between the three hydrogels with similar underlying crosslinking mechanisms but different end-point functionalities, we examined the generation of the proliferative phenotype of germinal center-like B cells, i.e. CD19+GL7+Fas+ cells. The PEG-4MAL organoids generated significantly higher number CD19+GL7+Fas+ B cells (Figure 2I), in contrast to PEG-4VS and PEG-4Acr. The Fas (CD95, Apo-1) receptor induces apoptosis in germinal centers upon engagement by Fas ligand (FasL) present on T_FH_ cells, and is one of many B cell proliferation restraining mechanism in vivo^28, 29^. Absence of Fas in B cell-specific Fas-deficient mice can result in fatal lymphoproliferation. Since our organoid system does not have a FasL checkpoint, we hypothesized that the polymer chemistry that supported induction of CD19+GL7+Fas+ B cells would allow for proliferation of germinal center-like B cells, and therefore an increased number of these cells should be present in the absence of a Fas-FasL check. To test this hypothesis, we examined the expression of Fas in the few surviving CD19+GL7+Fas+ B cells in PEG-4VS and PEG-4Acr, as compared to a large number of surviving CD19+GL7+Fas+ B cells in PEG-4MAL. The expression level of Fas (mean fluorescence intensity) was comparable in the surviving CD19+GL7+Fas+ B cells in PEG-4VS and PEG-4Acr as compared to PEG-4MAL (Figure 2J), creating a possibility that the ones that survive express high levels of Fas and plausibly the ones that die in PEG-4VS and PEG-4Acr had failed to express key proliferative signals. Further temporal analysis would be needed to understand the mechanism of Fas expression in dying cells in the context of polymer chemistry.

### 2.3. Collagen 1 mimicking peptide regulates germinal center phenotype and BCL6 expression towards germinal center exit-like phenotype

As indicated in Figure 1, various follicle-associated stromal cells, including FDCs, express the collagen 1 gene. This is further supported by the presence of collagen conduits in B cell follicles and whole lymph node^30^. How collagen-mimicking matrices compare to fibronectin, vitronectin, or VCAM-1 ligands in inducing germinal centers remains unknown. Here, we examined three synthetic adhesive peptides with different integrin binding specificities (Figure 3A). GFOGER is a triple helical synthetic peptide derived from type I collagen with high binding affinity for α1β1, α2β1, α10β1 and α11β1^31, 32^. REDV is a tetrapeptide Arg-Glu-Asp-Val that mimics VCAM-1 in its ability to bind α4β1 integrins^33^ on B cells, and RGD is a short linear peptide present in vitronectin, fibronectin and other ECM proteins that binds several integrins, including αvβ3, αvβ1, and α5β1^34–36^. We have previously shown that scrambled inactive peptides, such as G<u>RDG</u>SPC, do not engage integrins on B cells and fail to impact B cell receptors that are critical to downstream signaling and immune response ^20^, and therefore RDG only groups were not included in the current study. We did, however, incorporate RDG scramble peptides in the organoids as a filler to overcome solubility and viscosity issues when 100% GFOGER is used. Therefore, we used 0.3 molar ratio of GFOGER, REDV, or RGD and 0.7 molar ratio of RDG.

**Figure 3:**
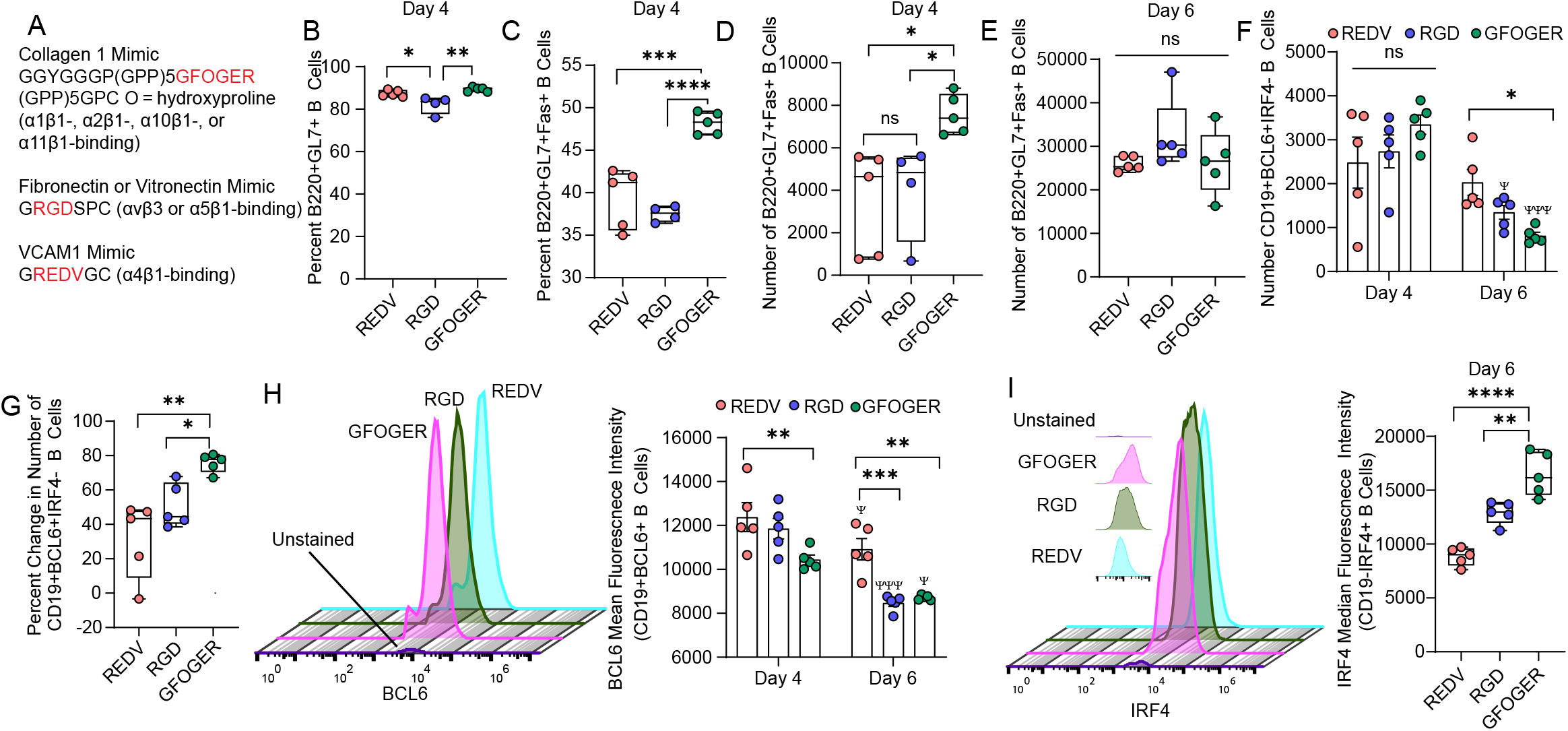
Collagen-mimicking GFOGER peptide ligand differentially regulates germinal center-like B cell activation and signaling. **A)** Sequences of integrin-binding peptides incorporated into PEG-4MAL organoids. **B-C)** Effects of GFOGER peptide ligand on percent of **B)** B220+GL7+ B cells and **C)** proliferative B220+GL7+Fas+ B cells, in contrast to VCAM-1-like and RGD peptide ligands in organoid grown cultures of naïve B cells with 40LB and IL-4. **D-E)** Time-dependent effects of varying integrin ligand on number of B220+GL7+Fas+ cells on **D)** Day 4 and **E)** Day 6. **F-G)** Temporal effect of integrin-binding peptide on **F)** number of and **G)** percent change in CD19+BCL6+IRF4-B cells over time. **H)** BCL6 mean fluorescence intensity over time in response to varying integrin ligand. Left side flow cytometry histogram indicates expression of BCL6 on Day 4. **I)** IRF4 mean fluorescence intensity over time in response to varying integrin ligand. Left side flow cytometry histogram indicates expression of IRF4 on Day 6. Data represent mean ± S.E.M. Significance determined via one-way ANOVA with Tukey’s post-hoc multiple comparisons test. *p<0.05, **p<0.01, ***p<0.001, ****p<0.0001; N=5.

While all three peptides supported induction of nearly similar percentage of CD19+GL7+ germinal center phenotype cells by day 4 in presence of 40LB cells (Figure 3B), GFOGER peptide functionalized PEG-4MAL organoids significantly increased the hallmark proliferative CD19+GL7+Fas+ phenotype by ~ 10% as compared to REDV and RGD peptides (Figure 3C). There were no significant differences between REDV and RGD peptides in the induction of CD19+GL7+Fas+ population. Similarly, we observed a significant increase in the number of CD19+GL7+Fas+ cells in GFOGER-organoids than in other matrices (Figure 3D). By day 6, the differences between matrices were lost, suggesting that matrices can play a role in the initial time course of induction of the germinal center phenotype (Figure 3E). Such studies are difficult to perform *in vivo* due to the migratory behavior of B cells that manifest unique cell cycle stages and emphasize the need for *ex vivo* immune organoids where nearly all cells are synchronized to similar states. By day 6, the expression of GL7 significantly reduced compared to day 4 (**Supplementary Fig 1A)**. There were no differences across matrices on day 6, possibly suggesting further differentiation of germinal center-like B cells towards a germinal center exit phenotype.

We next studied the temporal expression of hallmark transcription factor, BCL6, in germinal center-like B cells differentiated in organoids. At the transcriptional level, maintenance of the germinal center B cell fate is regulated by germinal center promoting transcription factors, in particular Bcl6, and B cell exit to plasma cell fate is controlled by transcription factors such as Irf4 ^37, 38^. Towards the exit of germinal centers, plasma-cell-precursor-like cells express lower Bcl6 ^39^. There were no differences in the number of CD19+BCL6+IRF4-B cells on day 4 of differentiation across the three matrices (Figure 3F). In contrast, the number of CD19+BCL6+IRF4-B cells reduced by day 6 (Figure 3F), with a decrease by 31 ± 22 % in REDV, 50 ± 12 % in RGD, and maximum reduction in GFOGER-organoids with a drop by 76 ± 5 % (Figure 3G). The decrease in CD19+BCL6+IRF4-cells does not indicate a decrease in differentiated B cells because there are > 90% B220+GL7+ B cells on day 6 **(Supplementary Fig 1B)**. It could reflect that the activated B cells that acquired a GL7 phenotype are differentiating into an advanced stage by day 6, possibly progressing towards a germinal center exit phenotype *ex vivo* that has a marked decrease in BCL6 expression. To test this, we evaluated the expression of BCL6 in CD19+BCL6+ B cells and observed a significant decrease in BCL6 expression on day 6 and in a matrix-dependent manner with cells grown in GFOGER and RGD organoids acquiring more reduction than REDV organoids (Figure 3H). These results suggest that GFOGER-presenting immune organoids induce high differentiation of germinal center B cells within 4 days of differentiation with rapid transition to germinal center exit-like phenotype by day 6. Our hypothesis is further supported by higher expression of IRF4 in B220-IRF4+ B cells in the GFOGER-functionalized organoids as compared to REDV- or RGD-functionalized organoids (Figure 3I). IRF4 is a transcription factor that is obligatory required for terminal differentiation of B cells to plasma cells^38^.

GFOGER-like sequences in collagens can bind to α1β1, α2β1, α10β1 and α11β1 integrins with high affinity. Our RNA sequencing results suggest that both naïve and in vivo germinal center B cells lack α1, α2, α10 and α11 integrins, however express high levels of β1 integrins. Yet, other studies have shown that integrins α1β1 and α2β1 are present on activated T-cells and fibroblasts (α1β1, α2β1), among other cell types ^32^. In addition, integrin α11β1 is also present on a subset of fibroblasts ^32^. Because 40LBs are fibroblasts, it is unclear whether GFOGER binds directly to B cells in organoids or functions via 40LB cells, or whether the effect is mediated by high expression of β1 and its interaction with the triple helical GFOGER peptide as compared to the linear REDV and RGD peptides. Therefore, further investigation is needed to determine the mode of functioning of GFOGER in organoids.

### 2.4. GFOGER and REDV regulate germinal center B cell epigenetic hallmarks EZH2 and H3k27Me3 in a CD40L- and BCL6-dependent manner

Germinal center B cells feature upregulation of Polycomb protein Enhancer of zeste homolog 2 (EZH2) histone methyltransferase^19, 40^, a core component of Polycomb repressive complex (PRC) 2 that methylates lysine 27 of histone 3 to generate H3K27me3, a histone mark associated with gene repression. EZH2 converts H3K4me3 active promoters to H3K4me3/H3K27me3 bivalent promoters for transient repression of target genes, which is reversed upon germinal center exit. We and others have previously shown that EZH2 is required for germinal center formation and conditional deletion of EZH2 results in failure to form germinal centers in mice^19, 41^. We have also shown that germinal center B cells undergo massive proliferation due to silencing of cell cycle checkpoint genes, such as CDKN1A ^19^, at the epigenetic level, mediated by EZH2, which catalyzes H3k27Me3. EZH2 is absent from naïve B cells but highly induced in germinal centers. While it is known that α4β1-integrin mediated T cell adhesion to VCAM-1 drives histone 3 methylation through a methyltransferase, the role of various integrin ligands, including collagen 1, in modulating the EZH2 and H3k27Me3 kinetics of the germinal center differentiation remains unknown. Unfortunately, the crucial EZH2 kinetics experiments cannot be easily performed *in vivo* because: A) in mice, germinal centers are heterogeneous with many B cells entering, leaving and recycling at any given time, and B) knockdown of integrins and their ligands can be lethal. Developing biomaterials to recapitulate the process of germinal centers could improve the mechanistic investigation into signaling and epigenetic mechanisms that regulate germinal center B cells.

Along these lines, we had previously reported a gelatin-silicate nanoparticle-based immune organoid that presented RGD motifs to B cells ^19, 42^. In these studies, the immune organoids showed remarkable similarity to *in vivo* germinal center B cells transcriptome, somatic hypermutation, EZH2 and H3k27Me3 protein expression, and proliferative phenotype. Analysis of the *EZH2* gene from RNA sequencing results show that RGD organoids with primary B cells, 40LBs, and IL-4 manifest similar *EZH2* upregulation on day 4 as immunized mice on day 10 (typical time for formation of *in vivo* germinal centers) (Figure 4A). In contrast, 2D cocultures of primary naïve B cells and 40LBs in the presence of IL-4 failed to induce expression of *EZH2* and were similar to the levels in naïve B cells, which do not have high expression of EZH2 (Figure 4A). To determine whether REDV and GFOGER induced differential expression of EZH2 than in RGD, we evaluated the impact of these peptides on germinal center-like B cells differentiated in PEG-4MAL organoids.

**Figure 4:**
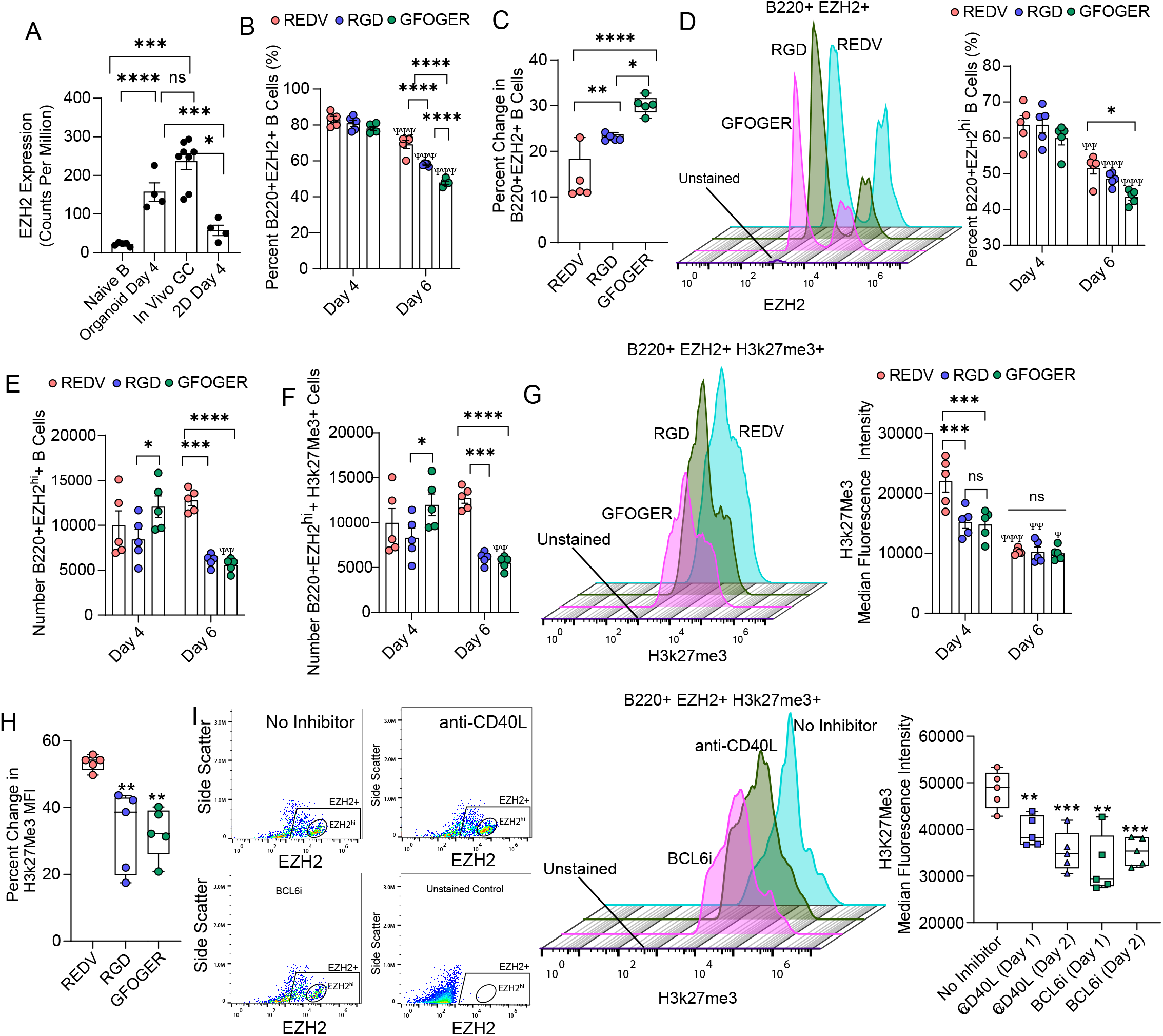
Expression of germinal center-like B cell epigenetics is regulated by Maleimide-presented integrin ligands. **A)** EZH2 expression in naïve murine B cells, generation 1, ex vivo 3D organoids made of RGD-rich matrix, in vivo germinal centers in immunized mice, and ex vivo 2D co-cultures. **B)** Effect of integrin-binding peptides on percentage of B220+EZH2+ B cells over time in organoid grown cultures of naïve B cells with 40LB and IL-4. **C)** Change in B220+EZH2+ B cells between days 4 and 6. **D-E)** Effect of integrin-binding peptides on **D)** percentage and **E)** number of B220+ B cells with high EZH2 expression. Left side flow cytometry histogram in D) indicates expression of EZH2 on Day 6. **F-G)** Effect of integrin-binding peptides on histone methylation in terms on **F)** number of positive cells and **G)** median fluorescence intensity. Left side flow cytometry histogram in G) indicates expression of H3k27Me3 on Day 6. **H)** Percent change in H3k27 methylation over time in response to varying integrin ligand. **I)** Effect of blocking CD40L and BCL6 on days 1 or 2 of ex vivo organoid culture on H3k27 methylation. Left side flow cytometry scatter plot and histogram indicates EZH2 populations and expression of H3k27Me3. Data represent mean ± S.E.M. Significance determined via one- or two-way ANOVA with Tukey’s post-hoc multiple comparisons test; N=5. *p<0.05, **p<0.01, ***p<0.001, ****p<0.0001; Ψp<0.05, ΨΨp<0.01, ΨΨΨp<0.001, ΨΨΨΨp<0.0001 relative to day 4 for corresponding ligand.

We observed no significant difference in B220+EZH2+ B cells on day 4 across the matrices, however the EZH2+ cells decreased in a matrix dependent manner by day 6 (Figure 4B). Although the decrease in all three matrices was significant compared to day 4, REDV only reduced the cells by 14% and RGD by 24%, as compared to 30% reduction in EZH2 in GFOGER-organoids (Figure 4C). Notably, ~60-65% of the B220+ B cells were positive for EZH2^hi^ on day 4. Percent and number of B220+EZH2hi cells followed a similar trend with matrices and temporal response, and the number of cells did not change for REDV and RGD over 6 days but only for GFOGER-functionalized organoids (Figure 4D, E). Similarly, the number of B220+EZH2+H3k27Me3+ cells did not change for REDV and RGD but only for GFOGER-functionalized organoids. Nearly all EZH2hi+ cells were also positive for H3k27Me3 (Figure 4F). We next evaluated the expression of H3k27Me3, as it represents the methyltransferase activity of EZH2. On day 4, VCAM-1 mimicking REDV-organoids expressed the highest levels of H3k27Me3 (Figure 4G). In contrast, differences between ECMs were insignificant on day 6. The drop in H3k27Me3 expression by day 6 was significant across all matrices with ~53% drop in REDV compared to a significantly lower 32% drop in GFOGER and RGD-organoids (Figure 4H).

Finally, we sought to determine whether the change in H3k27Me3 expression was regulated by the presence of T cell signal CD40L and BCL6. To understand this, we inhibited the CD40-CD40L interaction by initiating the germinal center response and then inhibiting on day 1 or day 2. Likewise, we inhibited BCL6 expression on day 1 or day 2 of culture. After inhibition, on day 4, we observed a significant decrease in the expression of H3k27Me3 (Figure 4I), clearly indicating that the H3k27Me3 was regulated by the induction of the germinal center response in organoids.

### 2.5. Effect of bacterial antigen recognition on germinal center B cell differentiation in young mice

The rise of antibiotic resistance among clinically relevant bacteria is one of the most prominent public health challenges of the modern era. *Klebsiella pneumonia*, in particular, is one of the most frequent causes of nosocomial infections that causes many severe diseases including pneumonia, bacteremia, and urinary tract infection, and some recent isolates have developed pan-drug resistance. Among those at risk are infants (who have immature immune systems) and the elderly (who have waning immune defenses); both are subject to heightened incidence and severity of infections^43, 44^. Carbapenem-resistant *K. pneumoniae* infections have an approximate 42% mortality rate and require intensive antibiotic cocktails of last-resort antibiotics with toxic side effects. With options for post-exposure antibiotic treatment dwindling, there is an urgent need for alternative therapies against antibiotic-resistant bacteria, such as *K. pneumoniae*, to prevent infection and treat infected patients. The external leaflet of the outer membrane of *K. pneumoniae* is formed by lipopolysaccharide (LPS), a glycolipid composed of lipid A, core oligosaccharide and O-antigen that serves as a defense barrier in bacteria. Recently, O-antigen-specific monoclonal antibodies against *K. pneumoniae* surface polysaccharides have been characterized as pre-emptive therapy but are still limited to a narrow spectrum of serotypes, necessitating the need to develop more of such advanced therapeutics.

As a classic T cell–independent antigen, *K. pneumoniae’s* glycan antigens fail to induce strong germinal center reactions to generate and select high-affinity antibody variants by affinity maturation. In contrast, T cell–dependent responses are typically directed against protein antigens and although *K. pneumoniae* harbors several surface proteins that could potentially serve as germinal center inducing agents, these are not well studied in the context of humoral immunity. A potential solution is to identify antigens and induce ex vivo germinal center responses to generate antibody secreting cells and subsequently, antibodies against these antigens. However, it remains unclear whether such antigenic proteins from Klebsiella would need to be presented as a soluble antigen or embedded within the bacterial membrane.

Here, as a proof of concept, we attempted to understand the latter question by testing the ability of *K. pneumoniae* membrane embedded OmpA protein (KlebOM) versus soluble, purified OmpA protein (sOmpA) in inducing *ex vivo* germinal center responses in PEG-4MAL organoids (Figure 5A). The use of immune organoids allows us to bypass the need for these antigens to induce CD40L expression and differentiation of T_FH_ cells *in vivo*, and yet test the hypothesis that membrane embedded protein antigens may induce stronger or weaker germinal center responses than soluble antigens (both antigen forms are expected to bind to the B cell receptor). Importantly, immune organoids provide a useful platform to potentially generate *ex vivo* antibodies against strains of bacteria. This proof of concept study focuses on the former question to optimize the mode of presentation in immune organoids.

**Figure 5:**
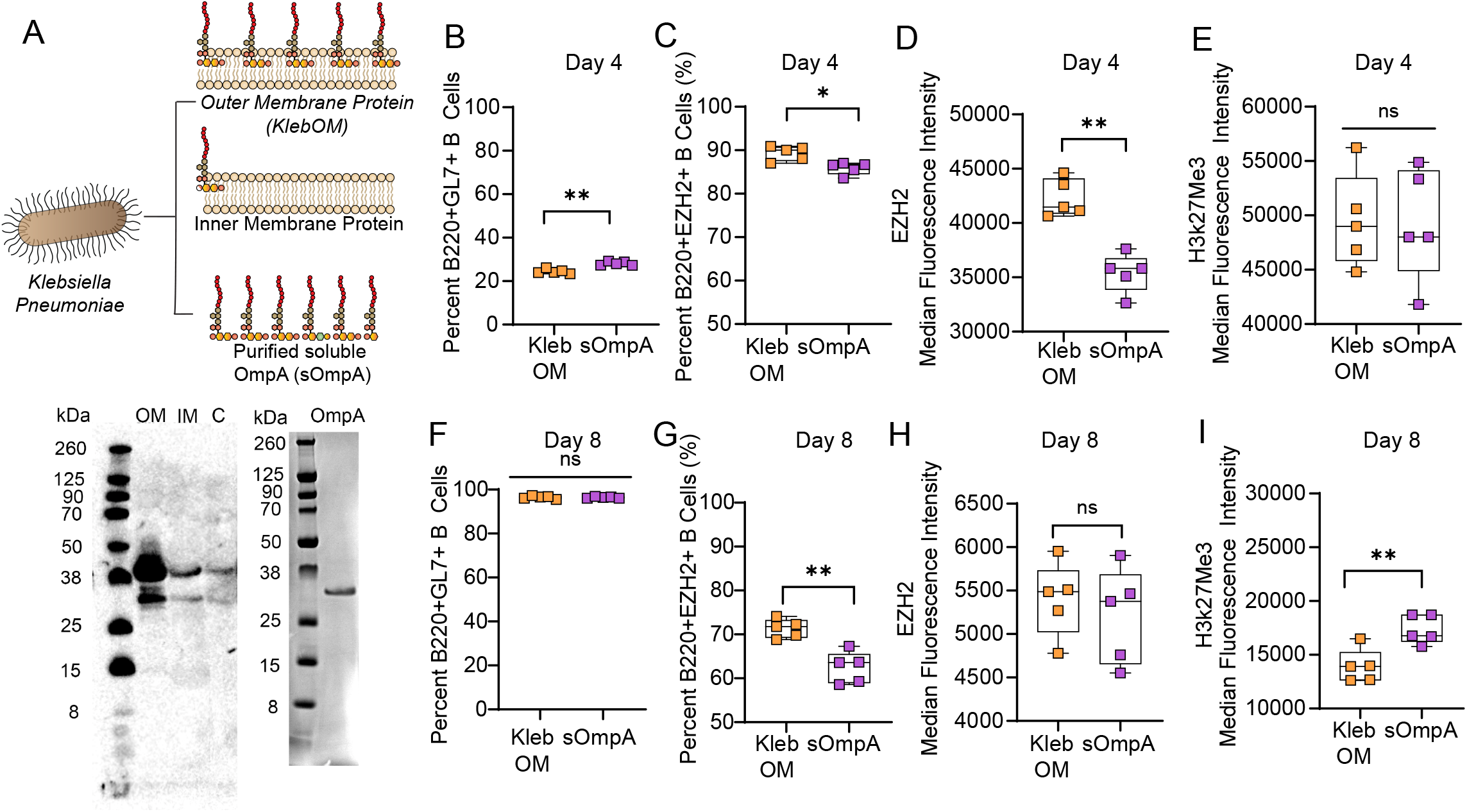
Mode of bacterial antigen presentation from clinical isolate of *Klebsiella pneumonia* determines the fate of germinal center B cell differentiation in young mice. **A)** Schematic of bacterial outer membrane-bound antigen (KlebOM) derived from *Klebsiella pneumonia* and soluble OmpA (sOmpA) protein extracted from outer membrane. Western blot showing the band of an outer membrane protein in the total outer membrane fraction, identified using an anti OmpA antibody. Adjacent western blot image shows the band of purified OmpA protein. OM: outer membrane, IM: inner membrane, C: control flow through. **B-E)** B cell response to membrane-embedded and soluble OmpA antigen in terms of **B)** percent B220+GL7+ cells, **C)** percent B220+EZH2+ cells, **D)** intensity of EZH2 expression, and **E)** intensity of histone methylation on day 4 of ex vivo PEG-4MAL organoid culture with 40LB and IL-4. **F-I)** B cell response to membrane-bound and soluble antigen in terms of **F)** percent B220+GL7+ cells, **G)** percent B220+EZH2+ cells, H) intensity of EZH2 expression, and **I)** intensity of histone methylation on day 8 of ex vivo organoid culture. Data represent mean ± S.E.M. Significance determined via unpaired, non-parametric Mann Whitney t-test; N=5. *p<0.05, **p<0.01, ns denotes no significance.

We added approximately 3.2 μg sOmpA or KlebOM to the media for each organoid. The soluble protein antigen, sOmpA, of *K. pneumoniae* induced modestly higher B220+GL7+ B cells than membrane embedded OmpA, KlebOM, which also contains other membrane bound components such as LPS (Figure 5B). Notably, the overall percentage of B220+GL7+ B cells were markedly low (~20-30%), which is possibly attributable to the very nature of the material composition of *K. pneumoniae* that induces poor germinal center response *in vivo*. Nearly 90% of these B220+GL7+ B cells expressed EZH2 with significantly higher EZH2 expressed by cells exposed to KlebOM than sOmpA (Figure 5C, **D**). Interestingly, the differences did not lead to change in the expression of H3k27Me3 by day 4 (Figure 5E). In contrast to day 4, by day 8, >90% cells became B220+GL7+, suggesting a delayed induction of germinal centers (Figure 5F). We chose day 8 to allow sufficient number of germinal center-like B cells in organoids. The percentage of B220+GL7+EZH2+ cells, however, declined to 60-75% (Figure 5E) and the differences in EZH2 expression became indistinguishable (Figure 5F). More importantly, the expression of H3k27Me3 by day 8 increased in organoids exposed to sOmpA. These findings suggest that, at least *ex vivo*, histone methylation is more robust with sOmpA than membrane embedded antigen, highlighting the potential for future purified protein antigens without added LPS components.

### 2.6. Organoids predict the efficacy of bacterial antigen recognition on germinal center B cell differentiation in aged mice

We next sought to investigate the efficacy of KlebOM and sOmpA on germinal center B cell differentiation in aged mice and determine whether antigen mediated immune response can be generated in B cells from aged individuals, who are at increased risk of *K. pneumoniae* infections^44^. Aging mice display many similar features of human immune aging - including reduced humoral immune responses to vaccination, which has been attributed to a defective T_FH_ cell function in germinal centers^45^ and immunosenescence of B cells including reduced expression of genes crucial to differentiation and somatic hypermutation^46^. However, the precise immunological mechanisms remain unclear and therefore a more reliable, higher throughput approach to generate antibodies is using organoids. We tested the ability of organoids to induce germinal center B cell responses in primary B cells derived from aged mice and, remarkably, we saw a reduction in the number of B220+GL7+ B cells, expression of EZH2, and number of B220+GL7+H3kme273+ B cells in organoids without any antigen simply under the effect of 40LBs and IL-4 (Figure 6A-C). An important observation in this study is that when aged B cells are provided similar levels of CD40L (i.e. T_FH_ signal), they still do not get activated and proliferate as much as B cells from young mice. Nevertheless, the organoids generated sufficient number of germinal center B cells to perform a comparison between KlebOM and sOmpA in an aged setting. While both antigen forms led to a similar percentage of GL7+ B cells, the proportion of B220+EZH2+ and H3k27Me3+ B cells were lower in sOmpA conditions (Figure 6E,F). Nevertheless, the expression of H3k27Me3 in B220+EZH2+H3k27Me3+ B cells were higher in conditions with sOmpA (Figure 6G) and correlated with a concomitant increase in BCL6 (Figure 6H). Therefore, it is evident that by using organoids, we can generate germinal center-like B cells in aged mice, that are initially low in numbers but follow a trajectory similar to young B cells in predicting the response of sOmpA versus KlebOM.

**Figure 6:**
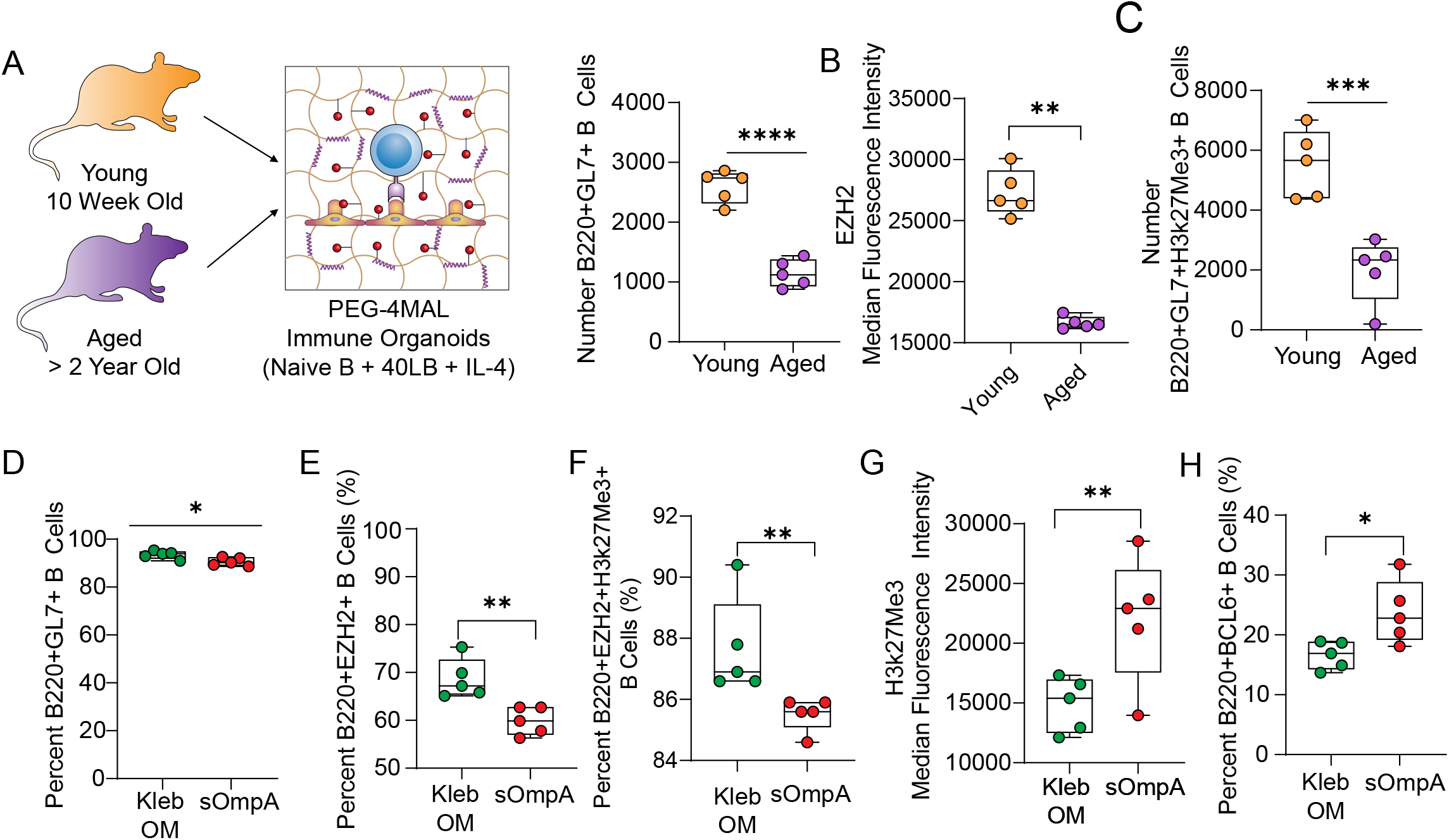
PEG-4MAL organoids induce germinal center reaction in aged B cells and predict immunogenic performance of Klebsiella pneumonia protein antigen. **A-C)** B cell response in the absence of any antigen in terms of A) number B220+GL7+ cells, **B)** intensity of EZH2 expression, and **C)** number of B220+EZH2+H3k27Me3+ B cells on day 4 of ex vivo PEG-4MAL organoid culture with 40LB and IL-4. **D-H)** Response to membrane-bound and soluble antigen in terms of **D)** percent B220+GL7+ cells, **E)** percent B220+EZH2+ cells, **F)** percent B220+EZH2+H3k27me3+ cells, **G) i**ntensity of histone methylation, and **H)** percent B220+BCL6+ cells on day 8 of ex vivo organoid culture using B cells derived from aged mice. Data represent mean ± S.E.M. Significance determined via unpaired, non-parametric Mann Whitney t-test; N=5. *p<0.05, **p<0.01, ns denotes no significance.

## 3. Conclusion

This work brings together several innovative concepts together, ranging from the effect of polymer end-chemistry on primary immune cells survival and differentiation, single cell and bulk RNA sequencing to define lymph node environment factors and the effect of these factors on the temporal kinetics of B cell differentiation, investigation of the mode of antigen formulation from clinical isolate of antibiotic-resistant *Klebsiella pneumoniae* on the temporal kinetics of B cell differentiation in young and aged mice, and the first demonstration of ex vivo B cell differentiation from primary immune cells derived from >2 year old aged mice. *Ex vivo* organoid studies offer a unique opportunity to identify protein antigens and conditions that may induce a more uniform response across both young and aged populations. This platform also offers an opportunity to derive age-specific germinal center B cells, and subsequently antibody secreting cells and antibodies, which can be injected into patients as therapeutics. This aspect may avoid a vaccine development that still requires induction of germinal centers *in vivo* and will, in the long term, provide alternative solutions to overcoming antibiotic-resistant and regular bacterial infections. Future work will focus on determining whether the two antigen forms can induce distinct or comparable somatic hypermutation and affinity maturation, generate protective antibodies against *K. pneumoniae* and be useful in aged mice with a defective T_FH_ cell function.

## 4. Materials and Methods

### Biomaterials and peptides

Four-arm polyethylene glycol-maleimide (PEG-MAL) with 20 kDa molecular weight and ≥90% purity was purchased from Laysan Bio. Integrin αvβ3-binding RGD peptide (G<u>RGD</u>SPC, >90% purity), scrambled peptide (G<u>RDG</u>SPC, >90% purity), integrin α4β1-binding REDV peptide (G<u>REDV</u>GC, >90% purity), GFOGER peptide (GYGGGPGPPGPPGPPGPPGPP<u>GFOGER</u>GPPGPPGPPGPPGPPGPC, ≥90% purity), and matrix metalloproteinase (MMP)-9 degradable VPM peptide (GCRD<u>VPM</u>SMRGGDRCG, >90% purity) were purchased from Aapptec. Non-degradable dithiothreitol (DTT) crosslinker was purchased from Sigma Aldrich. All components were reconstituted in 0.01 M HEPES buffer with pH 7.4.

### Stromal cell culture and mitomycin-c treatment

40LB stromal cells, genetically modified from NIH/3T3 fibroblasts to express CD40 ligand and produce B cell activating factor, as described previously ^19, 20, 25–27, 47^, were obtained from Dr. Daisuke Kitamura ^27^. 40LB cells were cultured in Dulbecco’s Modified Eagle Medium containing 10% fetal bovine serum and 1% penicillin streptomycin, and routinely passaged at 90% confluency. Prior to encapsulation in organoids, 40LB were mitotically inhibited via incubation with 0.01 mg/mL mitomycin-C at 37°C for 45 min.

### Naïve B cell isolation from mice

For experiments involving wildtype (WT) B cells, spleens were harvested from female C57BL/6 mice, aged 8-15 weeks, from the Jackson Laboratory. Where indicated, mice aged >2 years were used to compare the effects of aging. Spleens were dissociated using a sterile plunger, as previously described ^26^, and the cell suspension was incubated in RBC lysis buffer for 5 min at room temperature to remove red blood cells. Naïve B cells were purified from the splenocyte suspension by negative isolation using an EasySep Mouse B Cell Isolation Kit from Stem Cell Technologies, according to the manufacturer’s instructions. All animals were handled in compliance with the procedures approved by the Institutional Animal Care and Use Committee (IACUC) at Cornell University.

### Organoid fabrication

Synthetic immune organoids containing 7.5% PEG-4MAL or 7.5% PEG-4VS or 7.5% PEG-4Acr were fabricated using a 4-arm PEG-MAL macromer, adhesives peptides and crosslinkers. PEG-4MAL was functionalized at pH 7.4 with thiolated adhesive peptides RGD, REDV, or GFOGER at a 4:0.3 MAL-to-peptide molar ratio, and RDG scramble at a 4:0.7 MAL-to-peptide molar ratio for 30 min at 37°C. MMP9-degradable VPM peptide and non-degradable DTT crosslinkers were combined at a 1:1 VPM-to-DTT molar ratio and a 4:1.5 MAL-to-crosslinker molar ratio and adjusted to pH 5-6. Naïve B cells and 40LB stromal cells were suspended in the crosslinker solution, and 5 µL of crosslinker-cell suspension was injected into 5 µL functionalized PEG-MAL in each well of a non-treated 96 well plate. The droplet was mixed rapidly via pipet and cured at 37°C for 15 min. Post-incubation, Roswell Park Memorial Institute (RPMI) 1640 media containing 10% FBS, 1% P/S and 10 ng/mL interleukin-4 (IL-4, Peprotech) was added to each organoid. Media was replenished every 3 days.

### Inhibition of CD40L and BCL6

Where indicated, organoid media was replenished on days 1 or 2 with 50 μg/mL mouse anti-CD40L (Clone MR-1, Bio X Cell #BE0017-1), 50 μM BCL6 inhibitor 79-6 (Sigma #197345), or 10 ng/mL IL-4 alone. Changes in germinal center-like B cell dynamics were assessed on day 4 of organoid culture using flow cytometry.

### Enrichment of *K. pneumoniae* outer membranes (KlebOM)

Outer membranes were enriched from a clinical *Klebsiella pneumoniae* stool isolate, designated at B308-2, provided and profiled for antibiotic resistance by Michael Satlin at Weill Cornell Medicine. This isolate is here referred to as KpB308-2. KpB308-2 was inoculated in 500mL Nutrient Broth with 100ug/mL ampicillin and incubated overnight at 37°C, shaking 220 rpm. Following incubation, cells were pelleted at 5,000xg for 40 minutes at 4°C and resuspended in 36 mL lysis buffer (20mM Tris Buffer pH 8.0, 750mM Sucrose,) with 10 mg/mL egg white lysozyme and incubated on ice shaking 180 rpm for 1 hour. Ice cold 1.5mM EDTA was slowly poured into the lysozyme-treated cells and 1x protease inhibitor was added (Roche cat no. 04693159001). The suspensions were sonicated in an ice/water slurry for eight minutes total with the following parameters: 20 seconds on/20 seconds off, 70% amplitude. 1mM PMSF was added to the lysates, which were then spun at 1,500xg for 15 minutes at 4°C to remove large debris and unlysed cells. The supernatant was collected and spun at 100,000xg for 1 hour, 4°C to pellet membranes. The supernatant containing the soluble cytosolic fraction was removed and saved for analysis, and the pelleted membranes were resuspended in 2mL membrane wash buffer (20mM Tris Buffer pH 8.0, 300mM NaCl, 1x protease inhibitor, 1mM fresh PMSF) passaged through and 18g needle, washed in an additional 10mL supplemented wash buffer, and spun again at 100,00xg, 1 hour, 4°C. Total membrane pellets were resuspended via a 18g needle in 1% N-Lauroylsarcosine (Sigma #L5125) in supplemented wash buffer and incubated rolling at room temperature for 1 hour to selectively solubilize inner membranes ^48^. The remaining insoluble outer membrane fraction was pelleted at 100,000xg for 1 hour, 4°C. The supernatant containing solubilized inner membranes was removed and saved for analysis; and the enriched outer membrane pellet was resuspended once more in 10 mL supplemented membrane wash buffer, and pelleted a final time at 100,000xg, 4°C, for 1 hour. Finally, the enriched outer membrane pellet was resuspended in 3 mL of 20mM Tris-HCl, pH 8.0, 150mM NaCl, passed through a 0.22uM filter, and flash frozen until further use. Total protein concentration was measured using a Qubit 2. Approximately 3.2 μg KlebOM were supplemented to the media for each organoid.

### Western Blotting

To identify enrichment of KpB308-2 outer membranes, western blot was against the outer membrane marker Outer Membrane Protein A (OmpA). 15ug of total protein was mixed with 1X NuPage LDS sample buffer (NP0007) and 2.5% beta-mercaptoethanol, heated at 95°C for 5 minutes, then loaded into a 4-12% NuPage Bis-Tris gel. Protein was transferred to a nitrocellulose membrane using an iBlot 2 and blocked in 5% Skim milk powder in TBST for 1 hour at room temperature. The membrane was washed 3 times with TBST between each incubation. Membranes were incubated with a 1:5000 dilution of rabbit-anti-*Klebsiella* OmpA antibody (Antibody Research Corporation #111226) for 1 hour at room temperature, followed by incubation in a 1:10,000 dilution of HRP-conjugated goat-anti-rabbit secondary antibody (Thermo Fisher #65-6120). Blots were treated with Pierce ECL Western Blotting Substrate (Thermo Fisher #32106) and chemiluminescence was detected using the Azure Biosystems c300 Imaging system.

### Purification of *K. pneumoniae* Outer membrane protein A (sOmpA)

*K. pneumoniae* Outer membrane protein A (OmpA) was PCR amplified from genomic DNA from the clinical isolate B308-2 using the following primers: 5’ GATATACATATGAAAAAGACAGCTATCGCG 3’ and 5’ GTGGTGCTCGAGAGCCGCTGGCTGAGTTAC 3’ for insertion into the vector pET21b containing a C-terminal hexahistidine tag using the NdeI and XhoI restriction sites. Insertion was confirmed via Sanger sequencing using the primers: 5’ TAATACGACTCACTATAGGG 3’, 5’ GCTAGTTATTGCTCAGCGG 3’. The KpOmpA_pET21b construct was transformed into BL21 Star DE3 and plated on LB-Amp for overnight incubation at 37°C. Colonies were then scraped into 10 mL Terrific Broth with 100μg/mL ampicillin and grown for 2 hours before inoculating 1L Terrific Broth with 100μg/mL ampicillin. 1L cultures were grown shaking at 37°C to OD_600_ = 0.7 before adding 0.5mM IPTG to induce protein expression. Following IPTG addition, cultures were moved to shaking at 24°C overnight. After an overnight incubation, cultures were pelleted 5,000xg, 40 minutes, 4°C and resuspended in membrane wash buffer (20mM Tris Buffer pH 8.0, 300mM NaCl) and pelleted again. To lyse cells, the pellet was resuspended in 50 mL membrane wash buffer supplemented with 1X protease inhibitor (Roche cat no. 04693159001) and 10mg/mL lysozyme and let shake on ice for 30 minutes before sonication at 70% amplitude, 20 seconds on/ 20 seconds off for a total of 8 minutes. 1mM PMSF and 5mM Beta-mercaptoethanol were added to the lysates following sonication. Unlysed cells were removed via centrifugation at 1,500xg for 10 minutes, 4°C. Cleared lysates were spun at 100,000xg, 4°C for 1 hour, washed once with membrane wash buffer supplemented with 1x protease inhibitor and 1mM fresh PMSF, and spun again with the same settings. The pelleted total membrane fraction was resuspended once more in 3mL supplemented membrane wash buffer. To solubilize membranes and isolate protein, the resuspended membranes were mixed with 5% of 3:1 Styrene-maleic acid copolymer (3:1 SMA) in wash buffer and solubilized rolling at room temperature for two hours to allow the formation of membrane nanodiscs ^49^. Following incubation, insoluble material was pelleted at 10,000xg, 4°C, for 10 minutes and the nanodisc-solubilized membranes were mixed with 1mL of Qiagen Ni-NTA Superflow resin equilibrated with protein wash buffer (20mM Tris Buffer, pH 8.0, 500mM NaCl, 20mM Imidazole, 1x protease inhibitor, 10mM PMSF) was added to the cleared lysate and incubated rolling at room temperature for 1 hour to allow for protein binding. Lysates were loaded onto the column and washed with 40 resin volumes protein wash buffer prior to elution with 20mM Tris Buffer, 500mM EDTA, and 400mM Imidazole. Following elution, proteins were buffer exchanged and concentrated into 20mM Tris Buffer pH 8.0, 150mM NaCl using an Amicon 10 MWCO spin filter, passed through a 0.22uM filter to sterilize, and flash frozen until further use. 10 μL of purified protein was run on a 4-12% NuPage Bis-Tris gel with WesternSure Chemiluminescent protein ladder (Li-Cor 926-98000) and stained for total protein with SimplyBlue Safe Stain to confirm purity of sample. Approximately 3.2 μg sOmpA were supplemented to the media for each organoid.

### Flow cytometry analysis

On days 4 or 6, organoids were rinsed in 1X PBS and enzymatically digested in 125 U/mL collagenase type 1 (Worthington Biochemical) for 1 h at 37°C. Enzyme activity was terminated by addition of buffer containing serum, and the cells were filtered to remove organoid debris using 96 well MultiScreen Mesh Filter Plates (EMD Millipore). Cells were resuspended in FACS Buffer (PBS++ with 2% FBS, 1% P/S, and 0.005 mM EDTA) containing antibodies against cell surface antigens and incubated in the dark on ice for 1 h. Post-incubation, the cells were washed and resuspended in FACS Buffer. For intracellular antigens, the cells were fixed for 30 min in Fixation/Permeabilization Buffer (eBioscience Foxp3/Transcription Factor Staining Buffer) and incubated in permeabilization buffer containing antibodies for 1 h on ice, protected from light. Anti-mouse monoclonal antibodies targeting cell surface antigens included anti-GL7 (FITC, PE, eFluor660; clone GL 7; Thermo Fisher), anti-B220 (PE-Cy7; clone RA3-6B2; Thermo Fisher), anti-Fas (APC; clone 15A7; Thermo Fisher), anti-CD19 (FITC; clone 1D3; Thermo Fisher), anti-CD138 (FITC; clone 300506; Thermo Fisher), anti-IgD (PE; clone 11-26C; Thermo Fisher), and anti-CD38 (APC; clone 90; Thermo Fisher). Anti-mouse monoclonal antibodies targeting intracellular antigens included anti-H3k27me3 (AF488, AF647; clone C36B11; Cell Signaling), anti-EZH2 (eFluor 660; clone AC22; Thermo Fisher), anti-CBP (FITC, AF488; clone C-1; Santa Cruz), anti-H3k27ac (AF647; clone D5E4; Cell Signaling), anti-IRF4 (FITC, PE, PE-Cy7; clone 3E4; Thermo Fisher), and anti-BCL6 (PE, APC; clone BCL-DWN; Thermo Fisher).

### Statistical analysis

Statistical analysis was performed using GraphPad Prism software. Data analysis used an unpaired two-tailed t test or one-way analysis of variance (ANOVA) with Tukey’s post hoc test or two-way ANOVA with Sidak’s multiple comparison test. Quantitative analyses as scatter or bar graphs are presented as means ± SEM. In all studies, *P < 0.05, **P < 0.01, and ***P < 0.001 unless otherwise stated. Non-significance is denoted by “ns.”

## Supporting Information

Supporting Information is available from the Wiley Online Library or from the author.

## Acknowledgements

The authors acknowledge financial support from the National Institute of Allergy and Infectious Diseases of the US National Institutes of Health (5R01AI132738-03 awarded to A.S.), a US National Science Foundation CAREER award (DMR-1554275 awarded to A.S.), and the Innovative Molecular Analysis Technology program of the US National Cancer Institute (NIH R33-CA212968-01 awarded to A.S.). The authors acknowledge financial support from the Immunoengineering T32 training grant to S.P. (NIH and NIBIB, 1T32EB023860-01A1). Opinions, interpretations, conclusions, and recommendations are those of the authors and are not necessarily endorsed by the funding agency.

## Author Contributions

P.L.G. conducted all PEG-4MAL hydrogel studies, young and aged animal studies, bacterial antigen studies and analyzed data with A.S. K.L performed comparisons between hydrogel chemistries. J.C. provided single cell RNA sequencing on lymph node stromal cells and advice on germinal center biology. S.P and I.B. generated bacterial proteins, membranes, and western blots. A.S and P.G wrote the manuscript and all authors reviewed the manuscript and provided feedback. The concept was conceived by A.S. and funding was generated by A.S.

## Competing interests

The authors declare that they have no competing interests.

## Data and materials availability

All data needed to evaluate the conclusions in the paper are present in the paper and/or the Supplementary Materials. Single cell RNA sequence data have been deposited by Jason Cyster^50^ in GEO under ID code GSE112903. Bulk RNA sequencing on germinal center B cells and naïve B cells have been deposited in GEO with the primary accession codes GSE95491, as reported by us^19^.

---

**Supplementary Figure 1: Kinetics of GL7 expression in germinal center B cells in response to integrin ligand presentation. A)** Change in GL7 intensity over time in response to integrin-binding peptides. **B)** Effect of integrin-binding peptide on germinal center B cells (B220+GL7+) on day 6 of organoid culture. Data represent mean ± S.E.M. Significance determined via one-way (**A)** or two-way **(B)** ANOVA with Tukey’s post-hoc multiple comparisons test; N=5. *p<0.05, ***p<0.001, ns=not significant; ^ΨΨΨΨ^p<0.0001 relative to day 4.

**Supplementary Figure 2: Change in EZH2 expression in response to integrin ligand presentation.** Data represent mean ± S.E.M. Significance determined via one-way ANOVA with Tukey’s post-hoc multiple comparisons test; N=5 and ns denotes no significance.

## References

1. Guilbert, J.J. The World Health Report 2006: working together for health. Educ Health (Abingdon) 19, 385–387 (2006).

2. CDC. Antibiotic Resistance Threats in the United States, 2019. Atlanta, GA: U.S. Department of Health and Human Services, CDC; 2019. https://www.cdc.gov/drugresistance/Biggest-Threats.html. (2019).

3. McKenna, M. Antibiotic resistance: the last resort. Nature 499, 394–396 (2013).

4. Pennini, M.E. et al. Immune stealth-driven O2 serotype prevalence and potential for therapeutic antibodies against multidrug resistant Klebsiella pneumoniae. Nat Commun 8, 1991 (2017).

5. Rollenske, T. et al. Cross-specificity of protective human antibodies against Klebsiella pneumoniae LPS O-antigen. Nature immunology 19, 617–624 (2018).

6. Cyster, J.G. Shining a light on germinal center B cells. Cell 143, 503–505 (2010).

7. Allen, C.D. et al. Germinal center dark and light zone organization is mediated by CXCR4 and CXCR5. Nature immunology 5, 943–952 (2004).

8. Allen, C.D., Okada, T. & Cyster, J.G. Germinal-center organization and cellular dynamics. Immunity 27, 190–202 (2007).

9. Tas, J.M. et al. Visualizing antibody affinity maturation in germinal centers. Science 351, 1048–1054 (2016).

10. Mintz, M.A. et al. The HVEM-BTLA Axis Restrains T Cell Help to Germinal Center B Cells and Functions as a Cell-Extrinsic Suppressor in Lymphomagenesis. Immunity 51, 310–323 e317 (2019).

11. Phan, T.G., Grigorova, I., Okada, T. & Cyster, J.G. Subcapsular encounter and complement-dependent transport of immune complexes by lymph node B cells. Nature immunology 8, 992–1000 (2007).

12. Wang, X., Rodda, L.B., Bannard, O. & Cyster, J.G. Integrin-mediated interactions between B cells and follicular dendritic cells influence germinal center B cell fitness. J Immunol 192, 4601–4609 (2014).

13. Schiemann, B. et al. An essential role for BAFF in the normal development of B cells through a BCMA-independent pathway. Science 293, 2111–2114 (2001).

14. Heesters, B.A., Myers, R.C. & Carroll, M.C. Follicular dendritic cells: dynamic antigen libraries. Nature Reviews Immunology 14, 495–504 (2014).

15. Rodda, L.B. et al. Single-Cell RNA Sequencing of Lymph Node Stromal Cells Reveals Niche-Associated Heterogeneity. Immunity (2018).

16. Wang, X., Rodda, L.B., Bannard, O. & Cyster, J.G. Integrin-mediated interactions between B cells and follicular dendritic cells influence germinal center B cell fitness. The Journal of Immunology 192, 4601–4609 (2014).

17. Allen, C.D. & Cyster, J.G. Follicular dendritic cell networks of primary follicles and germinal centers: phenotype and function. Semin Immunol 20, 14–25 (2008).

18. Guo, M. et al. EZH2 Represses the B Cell Transcriptional Program and Regulates Antibody-Secreting Cell Metabolism and Antibody Production. J Immunol 200, 1039–1052 (2018).

19. Beguelin, W. et al. EZH2 enables germinal centre formation through epigenetic silencing of CDKN1A and an Rb-E2F1 feedback loop. Nat Commun 8, 877 (2017).

20. Purwada, A. et al. Ex vivo synthetic immune tissues with T cell signals for differentiating antigen-specific, high affinity germinal center B cells. Biomaterials 198, 27–36 (2019).

21. Phelps, E.A. et al. Maleimide cross-linked bioactive PEG hydrogel exhibits improved reaction kinetics and cross-linking for cell encapsulation and in situ delivery. Adv Mater 24, 64–70, 62 (2012).

22. Goldberg, M.S. Improving cancer immunotherapy through nanotechnology. Nat Rev Cancer 19, 587–602 (2019).

23. Kim, S., Shah, S.B., Graney, P. & Singh, A. Multiscale engineering of immune cells and lymphoid organs. Nature Reviews Materials 4, 355–378 (2019).

24. Polini, A. et al. Towards the development of human immune-system-on-a-chip platforms. Drug Discov Today 24, 517–525 (2019).

25. Purwada, A. et al. Ex vivo engineered immune organoids for controlled germinal center reactions. Biomaterials 63, 24–34 (2015).

26. Purwada, A. & Singh, A. Immuno-engineered Organoids for Regulating the Kinetics of B cell Development and Antibody Production. Nature Protocols 12, 168–182 (2017).

27. Nojima, T. et al. In-vitro derived germinal centre B cells differentially generate memory B or plasma cells in vivo. Nat Commun 2, 465 (2011).

28. Watanabe-Fukunaga, R., Brannan, C.I., Copeland, N.G., Jenkins, N.A. & Nagata, S. Lymphoproliferation disorder in mice explained by defects in Fas antigen that mediates apoptosis. Nature 356, 314–317 (1992).

29. Takahashi, T. et al. Generalized lymphoproliferative disease in mice, caused by a point mutation in the Fas ligand. Cell 76, 969–976 (1994).

30. Cyster, J.G. B cell follicles and antigen encounters of the third kind. Nature immunology 11, 989–996 (2010).

31. Knight, C.G. et al. Identification in collagen type I of an integrin alpha2 beta1-binding site containing an essential GER sequence. The Journal of biological chemistry 273, 33287–33294 (1998).

32. Zeltz, C. & Gullberg, D. The integrin-collagen connection--a glue for tissue repair? Journal of Cell Science 129, 653–664 (2016).

33. Massia, S.P. & Hubbell, J.A. Vascular endothelial cell adhesion and spreading promoted by the peptide REDV of the IIICS region of plasma fibronectin is mediated by integrin alpha 4 beta 1. The Journal of biological chemistry 267, 14019–14026 (1992).

34. Clark, A.Y. et al. Integrin-specific hydrogels modulate transplanted human bone marrow-derived mesenchymal stem cell survival, engraftment, and reparative activities. Nat Commun 11, 114 (2020).

35. Arnaout, M.A., Mahalingam, B. & Xiong, J.P. Integrin structure, allostery, and bidirectional signaling. Annu Rev Cell Dev Biol 21, 381–410 (2005).

36. Bellis, S.L. Advantages of RGD peptides for directing cell association with biomaterials. Biomaterials 32, 4205–4210 (2011).

37. Basso, K. & Dalla-Favera, R. Roles of BCL6 in normal and transformed germinal center B cells. Immunol Rev 247, 172–183 (2012).

38. Klein, U. et al. Transcription factor IRF4 controls plasma cell differentiation and class-switch recombination. Nature immunology 7, 773–782 (2006).

39. Li, X. et al. Cbl Ubiquitin Ligases Control B Cell Exit from the Germinal-Center Reaction. Immunity 48, 530–541 e536 (2018).

40. Raaphorst, F.M. et al. Cutting edge: polycomb gene expression patterns reflect distinct B cell differentiation stages in human germinal centers. J Immunol 164, 1–4 (2000).

41. Béguelin, W. et al. EZH2 is required for germinal center formation and somatic EZH2 mutations promote lymphoid transformation. Cancer cell 23, 677–692 (2013).

42. Purwada, A. & Singh, A. Immuno-engineered organoids for regulating the kinetics of B-cell development and antibody production. Nat Protoc 12, 168–182 (2017).

43. Gorrie, C.L. et al. Antimicrobial-Resistant Klebsiella pneumoniae Carriage and Infection in Specialized Geriatric Care Wards Linked to Acquisition in the Referring Hospital. Clin Infect Dis 67, 161–170 (2018).

44. Podschun, R. & Ullmann, U. Klebsiella spp. as nosocomial pathogens: epidemiology, taxonomy, typing methods, and pathogenicity factors. Clin Microbiol Rev 11, 589–603 (1998).

45. Sage, P.T., Tan, C.L., Freeman, G.J., Haigis, M. & Sharpe, A.H. Defective TFH Cell Function and Increased TFR Cells Contribute to Defective Antibody Production in Aging. Cell reports 12, 163–171 (2015).

46. Cancro, M.P. et al. B cells and aging: molecules and mechanisms. Trends Immunol 30, 313–318 (2009).

47. Purwada, A., Shah, S.B., Beguelin, W., Melnick, A. & Singh, A. Modular Immune Organoids with Integrin Ligand Specificity Differentially Regulate Ex vivo B Cell Activation. ACS Biomater. Sci. Eng. 3, 214–225 (2017).

48. Hobb, R.I., Fields, J.A., Burns, C.M. & Thompson, S.A. Evaluation of procedures for outer membrane isolation from Campylobacter jejuni. Microbiology 155, 979–988 (2009).

49. Lee, S.C. et al. A method for detergent-free isolation of membrane proteins in their local lipid environment. Nature Protocols 11, 1149–1162 (2016).

50. Rodda, L.B. et al. Single-Cell RNA Sequencing of Lymph Node Stromal Cells Reveals Niche-Associated Heterogeneity. Immunity 48, 1014–1028 e1016 (2018).

