## Supplementary Figures for "Organoid Polymer Functionality and Mode of *Klebsiella Pneumoniae* Membrane Antigen Presentation Regulates Ex Vivo Germinal Center Epigenetics in Young and Aged B Cells"

### Supplementary Figure 1, Related to Figure 3

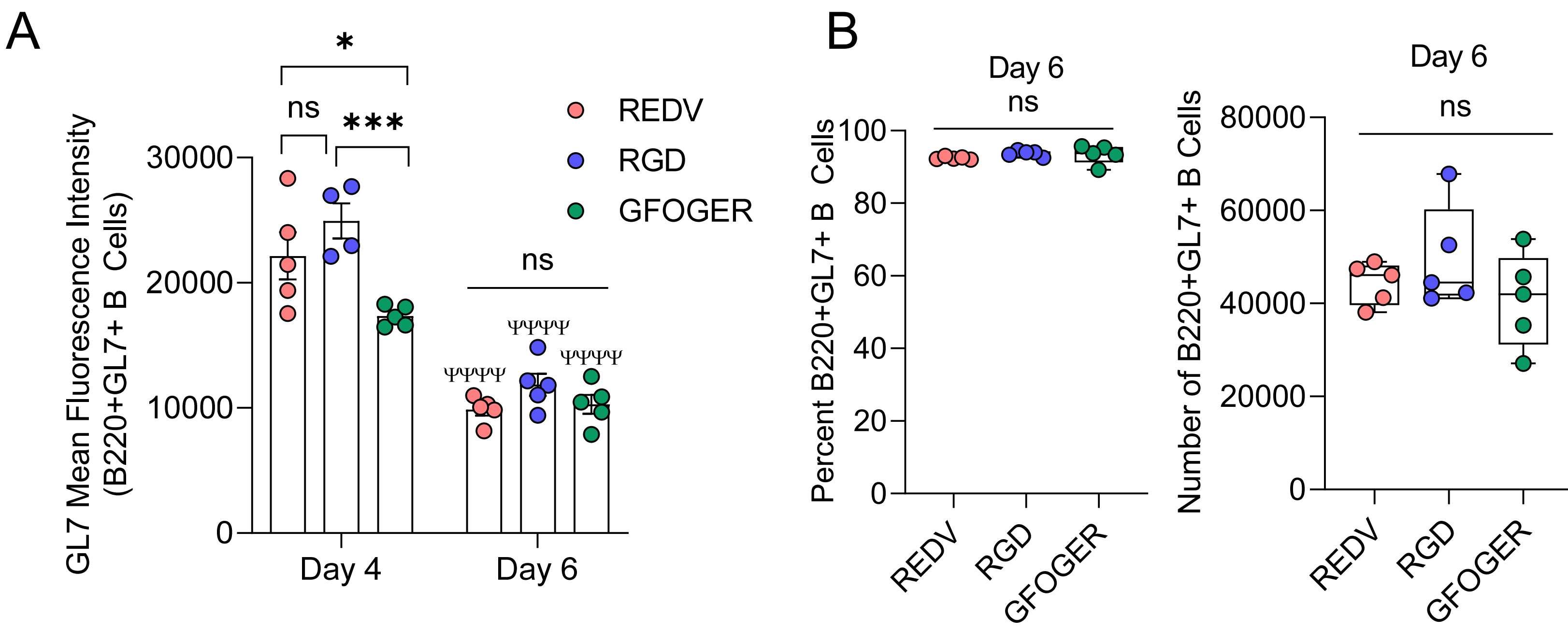


**Supplementary Figure 1: Kinetics of GL7 expression in germinal center B cells in response to integrin ligand presentation. A)** Change in GL7 intensity over time in response to integrin-binding peptides. **B)** Effect of integrin-binding peptide on germinal center B cells (B220+GL7+) on day 6 of organoid culture. Data represent mean ± S.E.M. Significance determined via one-way (A) or two-way (B) ANOVA with Tukey's post-hoc multiple comparisons test; N=5. *p<0.05, ***p<0.001, ns=not significant; ΨΨΨΨp<0.0001 relative to day 4.

Supplementary Figure 2, Related to Figure 3

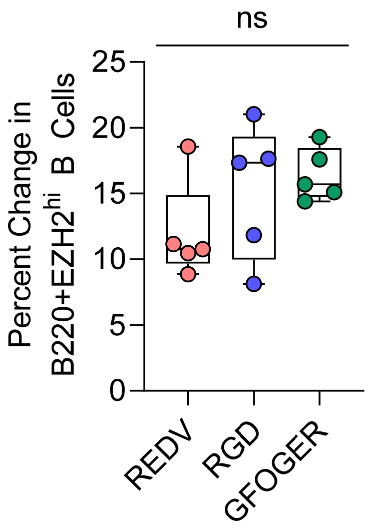


**Supplementary Figure 2: Change in EZH2 expression in response to integrin ligand presentation.** Data represent mean ± S.E.M. Significance determined via one-way ANOVA with Tukey's post-hoc multiple comparisons test; N=5 and ns denotes no significance.
